# Functional enrichments of disease variants across thousands of independent loci in eight diseases

**DOI:** 10.1101/048066

**Authors:** Abhishek K. Sarkar, Lucas D. Ward, Manolis Kellis

**Affiliations:** Computer Science and Artificial Intelligence Lab, Massachusetts Institute of Technology, Cambridge,MA, USA; The Broad Institute of MIT and Harvard, Cambridge, MA, USA

## Abstract

For most complex traits, known genetic associations only explain a small fraction of the narrow sense heritability prompting intense debate on the genetic basis of complex traits. Joint analysis of all common variants together explains much of this missing heritability and reveals that large numbers of weakly associated loci are enriched in regulatory regions, but fails to identify specific regions or biological pathways. Here, we use epigenomic annotations across 127 tissues and cell types to investigate weak regulatory associations, the specific enhancers they reside in, their downstream target genes, their upstream regulators, and the biological pathways they disrupt in eight common diseases. We show weak associations are significantly enriched in disease-relevant regulatory regions across thousands of independent loci. We develop methods to control for LD between weak associations and overlap between annotations. We show that weak non-coding associations are additionally enriched in relevant biological pathways implicating additional downstream target genes and upstream disease-specific master regulators. Our results can help guide the discovery of biologically meaningful, but currently undetectable regulatory loci underlying a number of common diseases.

## Introduction

Thousands of loci associated with hundreds of complex diseases have been reported in the NHGRI catalog of genome-wide association studies^1^ (GWASs). However, replicated genome-wide significant loci explain only a fraction of the heritability of complex traits, a discrepancy known as missing heritability, motivating inquiry into the architecture of complex disease^2^. Recent work modeling the joint effect of all SNPs supports a highly polygenic architecture^3,4^. For example, analysis of human height shows 16% of the phenotypic variance is explained by genome-wide significant loci, but 50% is explained by all SNPs^5^. Although this line of investigation has shed insight into complex diseases^6,7^ further work remains to identify the specific regions implicated.

Recent work also shows most genome-wide significant loci are devoid of protein-coding alterations^8^ and could instead affect transcriptional regulation. Associated loci are enriched in regulatory annotations including enhancers delineated by chromatin states^9^, DNaseI hypersensitive sites^10^ (DHSs), enhancer-associated histone modifications^11^, and large super-enhancers^12^. Moreover, enrichments persist beyond the traditional genome-wide significance threshold of *p* < 5 × 10^‒8^, providing a basis for re-prioritizing weak associations^13^.

Here we use regulatory annotations to go beyond identifying disease-relevant annotations by characterizing specific enhancer regions, their target genes, their upstream regulators, and the biological pathways disrupted by weakly associated non-coding variants. We combine and compare diverse regulatory annotations spanning multiple cell types, assays, and computational pipelines: chromatin states, DHSs, gene pathways, and regulatory motifs. We additionally control for a number of confounders including linkage disequilibrium (LD) between weak associations and overlap between regulatory annotations.

We carry out these studies in eight large scale meta-analyses of common diseases spanning autoimmune, psychiatric, and metabolic disorders. Across these eight diseases, we find thousands of independent loci are enriched for regulatory annotations in common pathways. We find enrichments for brain enhancers in bipolar disorder and schizophrenia; pancreatic islet enhancers in Type 2 Diabetes; mucosa enhancers in coronary artery disease; and immune enhancers in Type 1 Diabetes, Crohn’s disease, rheumatoid arthritis, and Alzheimer’s disease. We show regulatory variants disrupt both constitutive and tissue-specific enhancer regions predicted by chromatin marks. We find downstream target genes are enriched in a number of known biological pathways, but only a small fraction of the genes are already identified by GWAS. We identify upstream master regulators whose binding is indirectly disrupted and show that constitutively marked enhancer regions disrupted by weak associations may not be constitutively active due to tissue-specific expression of the upstream transcription factor. Together, our results illustrate an approach to identify many weakly associated common variants recurrently disrupting a small number of biological pathways in complex diseases.

## Results

### Functional enrichment of enhancer annotations

We investigated weak genetic associations (having *p* < 5.3 × 10^‒4^) with eight well-studied common diseases spanning a variety of etiologies, pathologies, and genetic architectures for which summary statistics are publicly available (**Supplementary Table 1**): Alzheimer’s disease (AD), bipolar disorder (BIP), coronary artery disease (CAD), Crohn’s disease (CD), rheumatoid arthritis (RA), schizophrenia (SCZ), Type 1 Diabetes (T1D), and Type 2 Diabetes (T2D).

The key idea of our approach is that the ranking of weak genetic associations gives partial information about the true underlying effects which can be used to identify enriched annotations. We first studied the robustness of the ranks to sample size using summary statistics for RA for which per-cohort *z*-scores and sample sizes were provided (**Supplementary Table 2**). We performed six meta-analyses holding out each cohort in turn and computed the correlation between *z*-scores in the held out cohort with the meta-analyzed *z*-scores of the remaining five cohorts. To account for inflation of test statistics around the Major Histocompatability Complex (MHC), we excluded chromosome 6. We also verified that the Pearson correlation was greater than 0.99 between our sample-size weighted meta-analyzed *z*-scores of the full study and the published inverse variance weighted *z*-scores. We found positive correlations between association *z*-scores on each cohort through tens of thousands of variants (assuming the original meta-analyzed *z*-scores are the true ranking) despite the fact that each individual cohort had between 483–1525 cases (**Supplementary Fig. 1**), supporting our idea that the ranking of *p*-values below genome-wide significance is informative of the ranking which would be obtained by a much larger study.

We next visualized enrichment of regulatory annotations using an approach inspired by Gene Set Enrichment Analysis^14^. Briefly, at every rank (*p*-value) threshold, we computed the difference between the observed number of overlaps with a regulatory annotation and the expected number, normalized by the total number of overlaps. Our visualization allows us to identify the relative importance of annotations based on the ordering of the curves and to determine an empirical *p*-value cutoff based on the elbow points of the curves.

We focus on distal enhancer regions because these play a role in transcriptional regulation and are also dynamic across different cell types, allowing us to propose causal cell types and tissue-specific biological functions which are disrupted. To define putative enhancer regions, we used a 15 chromatin state model^15^ summarizing five chromatin marks across 127 reference epigenomes spanning diverse primary cells and tissues from the Roadmap Epigenomics^16^ and ENCODE^17^ projects (**Supplementary Fig. 2**) and took the union of enhancer-like states.

We removed variants within the MHC (positions 29.4 – 33 MB of chromosome 6) plus 5 megabases flanking from all analyses. Although the MHC is known to play significant roles in diseases such as T1D, the causal variants in this region are known to be protein coding rather than regulatory, which is the focus of our study. Moreover, the MHC region displays unusual long range LD which inflates GWAS test statistics in the flanking regions and would confound our enrichments. In order to improve our power to detect enrichments, we imputed summary statistics for all studies into the Thousand Genomes reference cohort (if necessary) using ImpG-Summary^18^. In order to make our visualization comparable across different studies, we applied our method to a common set of 5.5 million well-imputed variants.

Applying our method to the eight diseases, we found enrichments for relevant cell types which persist even when considering thousands of weak associations, equal on average to a cutoff of *p* < 5.3 × 10^‒4^ (**Supplementary Fig. 3**). To account for linkage disequilibrium between weak associations, we adapted our visualization to work at the level of loci rather than variants, scoring each locus as the fraction of SNPs which have the annotation of interest. We pruned the imputed summary statistics to an average of 207,080 independent loci (pairwise *r*^2^ < 0.1) and found largely the same enrichments in each of the diseases through an average of 1,600 independent loci (**Fig. 1**).

**Figure 1:**
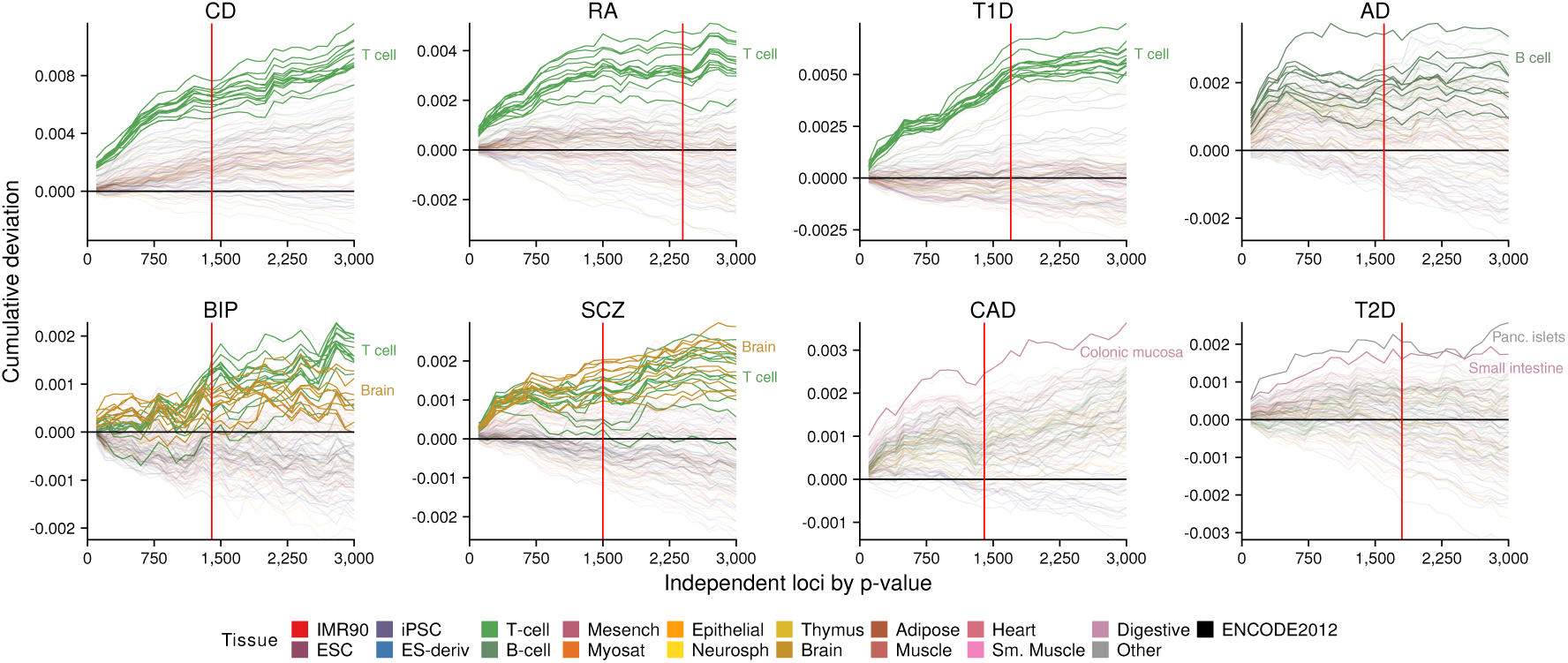
Enrichment of independent loci (pairwise *r*^2^ < 0.1) across eight diseases in enhancer regions predicted by a 15 chromatin state model learned on observed data for 5 histone modifications across 111 reference epigenomes. Each curve corresponds to enhancer regions predicted in a specific reference epigenome and is colored by tissue group. The black line at zero cumulative deviation indicates no enrichment, and the red vertical line indicates the empirical *p*-value cutoff taken forward for the rest of the analysis. *A priori* relevant enrichments are denoted by opaque lines.

In autoimmune disorders (CD, RA, T1D), we found enhancers active in T cell types showed the strongest enrichment for weak associations. In psychiatric disorders (AD, BIP, SCZ), we also found enrichment of immune cell types, supporting the role of immune pathways in these disorders^19–21^. Interestingly, we found enrichments for B cell types in addition to T cell types in BIP. In BIP and SCZ, we additionally found enrichment for enhancers in a number of adult brain tissues. In CAD, we found enrichments in colonic mucosa, which could be indicative of a role for endothelial cells for which epigenomic marks were not directly profiled. In T2D, we found enrichments in pancreatic islets, consistent with prior work^22^, but additionally in small intestine, consistent with the role of gastrointestinal mucosa in glucose homeostasis^23^.

We evaluated the statistical significance of enrichments using a permutation test based on Variant Set Enrichment^24^ (Online Methods). Briefly, for each disease and enhancer annotation, we compared the count of associations passing our empirical *p*-value cutoff within the annotation against the null distribution of counts of resampled SNPs passing the same cutoff outside the annotation. We resampled SNPs matched on LD block size, minor allele frequency, and distance to closest transcription start site. For each phenotype, we used all well-imputed SNPs (mean 7,797,600) to avoid small number effects. We found the enrichments reported above were all statistically significant after applying the Benjamini–Hochberg (BH) procedure with *q* = 0.05 (**Supplementary Fig. 4**); however, many additional cell types also showed significant enrichment attributable to confounding of constitutive and tissue-specific enhancers as we show below.

### Distinguishing constitutive and tissue-specific enhancers

We next sought to distinguish regions which exhibit enhancer-associated chromatin marks constitutively from those which are marked in specific tissues. Prior work has only considered annotations learned in individual cell types^25^, even when building joint models of a number of annotations. Here, we used 226 enhancer modules defined as previously described^16^ to delineate a biologically meaningful set of disjoint annotations. Briefly, putative enhancers across reference epigenomes are defined as DHSs (in any reference epigenome) labeled by enhancer-like chromatin states in each reference epigenome. Enhancer modules are then defined as *k*-means clusters of these regions based on their activity profiles across the reference epigenomes.

We computed enrichments for these enhancer modules and found that constitutive enhancers are significantly enriched for weak association across all eight diseases (permutation test, BH *q* = 0.05, **Fig. 2**). We note that these annotations cover such a small proportion of the genome that we could not use our visualization method to choose an empirical *p*-value cutoff, and instead used the cutoffs described above.

**Figure 2:**
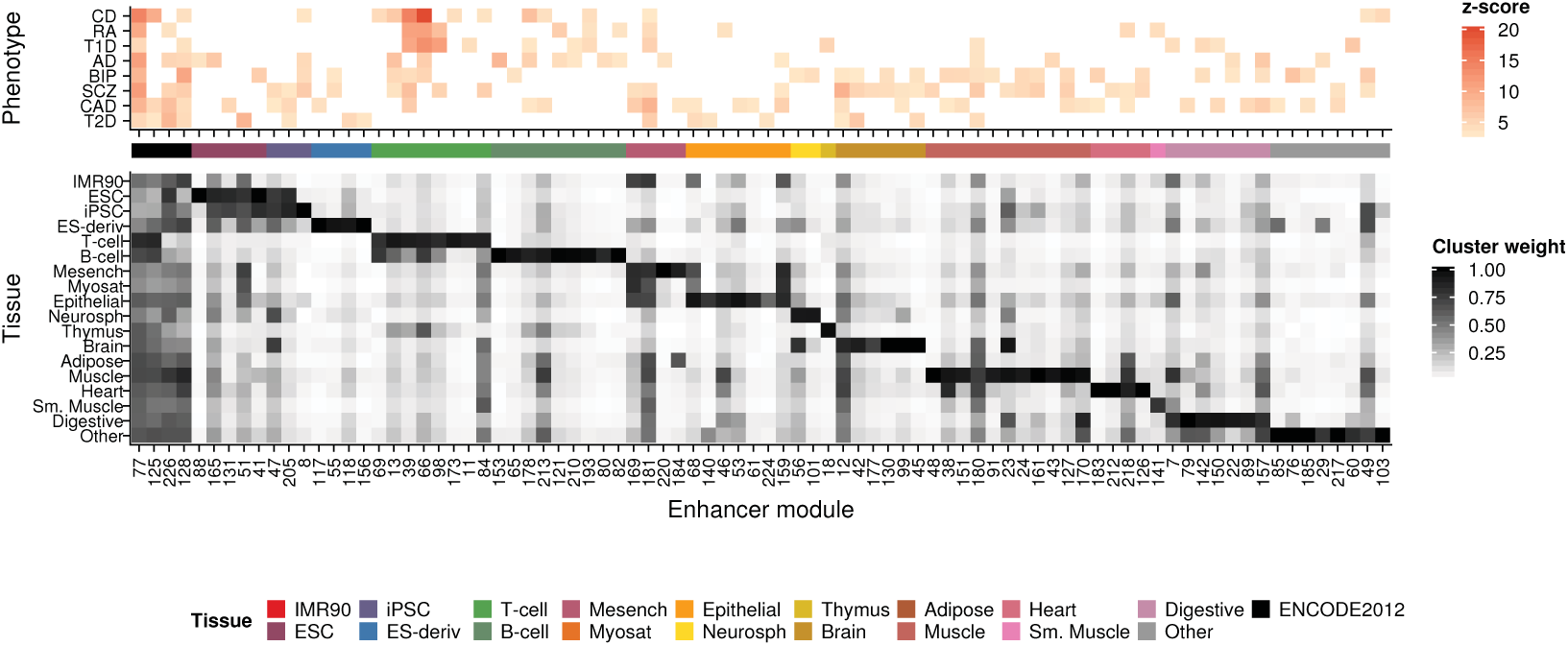
Enrichment of weak associations in enhancer modules. Enrichment *z*-scores for 226 enriched enhancer modules corresponding to observed histone modification patterns across 111 reference epigenomes. Only 84 significantly enriched modules (BH *q* < 0.05) are shown. Modules are defined by clustering DHSs labeled as enhancer-like by a 15 chromatin state model learned on observed data for 5 histone modifications across 111 reference epigenomes. Each module is represented by a vector of weights per reference epigenome (proportion of DHSs annotated as enhancer in that reference epigenome). For display, weights are collapsed by tissue group by taking the maximum weight over all reference epigenomes in each tissue group. Modules are ordered by the tissue group with maximum weight. The leftmost four modules are defined as constitutive (having at least 50% of cluster weights greater than 0.25).

After partitioning regulatory regions into constitutive and tissue-specific modules, we recover much fewer significant tissue-specific annotations. Our enrichments are less noisy not only because we correct for the contribution of constitutive enhancers to all single cell type annotations, but also because we use narrower, higher confidence regions by combining chromatin accessibility and histone modification data. We found that immune-specific enhancers are enriched in both autoimmune (CD, RA, T1D) and psychiatric disorders (AD, BIP, SCZ) and that brain-specific enhancers are enriched in psychiatric disorders. We found that mucosa-specific enhancers are enriched in metabolic disorders (CAD, T2D).

### Pathway enrichment of enhancer targets

We next investigated the target genes of enriched tissue-specific enhancer modules harboring weak associations. Prior work has used hierarchical modeling to study enrichment of weak associations in gene pathways^26^; however, current methods are limited to using proximity to link SNPs to their target genes, ignoring the regulatory potential of specific variants. We used GREAT^27^ to test genes linked to disrupted tissue-specific enhancers (as defined by the enriched modules above) for enrichment of Gene Ontology Biological Processes and took terms with FDR *q* < 0.05.

We found significant enrichments for a number of known pathways in each of the eight diseases (**Table 1**). In autoimmune disorders, we found enrichment for various pathways relating to immune response. However, we identified different specific signaling pathways in each disease: Immunoglobulin E and Interleukin-4 in CD, nuclear factor kappa-B in RA, and Interferon G in T1D. Surprisingly, we found enrichment for MHC class I/II processes in T1D despite excluding the MHC from the analysis. We verified this enrichment was not due to spurious correlations on chromosome 6. Instead, the enrichment is primarily driven by enhancers linked to *CIITA*, a known regulator of the MHC pathways which resides on chromosome 16.

**Table 1:** Pathway enrichments of enhancers harboring weak associations. Total gene counts are based on links to weakly associated enhancers across any significantly enriched pathway. Total pathway counts are restricted to GO terms with significant enrichments (FDR *q* < 0.05) for which no child (connected by an ontology relationship) is significantly enriched.

| Trait | Known pathways | Total genes | Total pathways |
| --- | --- | --- | --- |
| AD | Cyclic GMP signaling, immune response | 220 | 216 |
| BIP | Glucocorticoid signaling | 217 | 230 |
| CAD | Cholesterol/triglyceride metabolism, IgA | 248 | 215 |
| CD | CD8 T cell proliferation, IgE, IL4 | 224 | 359 |
| RA | NFKB, actin nucleation | 196 | 146 |
| SCZ | Dendritic spine development | 271 | 183 |
| T1D | MHC I/II, JAK-STAT, IFNG | 266 | 245 |
| T2D | Pancreatic $\beta$ cell apoptosis | 281 | 177 |

In psychiatric disorders, we recovered several known signaling pathways important to brain function such as cyclic GMP signaling in AD and glucocorticoid signaling in BIP, and brain development such as dendritic spine development in SCZ. We additionally found enrichment for immune response in AD, further supporting the role of immune pathways in this disease.

In CAD, we found enrichments for cholesterol and triglyceride metabolism, but additionally for the Immunoglobulin A pathway. In T2D, we found enrichment for pancreatic *β* cell apoptosis, a known hallmark of the disease.

We note that we recovered known pathways by considering weak associations which overlap distal regulatory regions rather than genome-wide significant associations which implicate nearby genes in LD. We used Phenotype-Genotype Integrator (PheGenI) to obtain lists of known genes for each disease and found that on average we linked putative disrupted enhancers to only 20 known genes across all enriched pathways for each disease (**Supplementary Table 3**). The remaining genes (**Supplementary Table 4**) are potentially new targets for experimental followup; however, we cannot assign a *p*-value to any particular gene.

Our approach yielded a large number of enriched GO terms and an average of 240 linked genes in each of the eight diseases. We used ontology relationships to prune the list of enriched terms to the most specific enriched terms. Briefly, we built a directed acyclic graph where nodes are GO terms and edges are ontology relationships and took all enriched nodes for which no child was enriched. Our approach recovered 146–359 enriched GO terms; however, we still recovered some *a priori* implausible pathways, possibly due to incorrect linking of enhancers to their target genes.

### Motif enrichment of upstream regulators

We next identified the upstream regulators whose binding may be perturbed by weak associations. Prior work has studied enrichment of regulatory motifs in enhancer regions^16,22^; however, these studies do not specifically consider the impact of SNPs on transcription factor binding affinity at specific motif instances. We studied regulatory motifs curated into 651 families^28^ and hypothesized that weak associations may recurrently affect binding of a small number of disease-specific master regulators by disrupting motif instances of co-factors^29^.

Briefly, we identified putative master regulators by studying the enrichment of motif instances in enhancer modules. We filtered motifs according to sequence enrichment against shuffled instances as previously described^28^. We then tested for enriched co-occurrence of weak associations and enriched motifs in each enhancer module using Fisher’s exact test. We finally re-scanned enhancer regions containing both a master regulator motif instance and a weak association to find co-occurring motifs which overlap weakly associated SNPs.

Our approach identified a number of significantly enriched master regulators across the eight diseases (Fisher’s exact test, BH *q* = 0.05, **Fig. 3**). Only three of the regulators have been previously identified by GWAS for the eight diseases and reported in PheGenI: *ETS1* in RA, *STAT3* in CD, and *NFKB1* in SCZ. This result is expected given that the majority of GWAS-identified loci do not implicate protein-coding genes; however, it also illustrates the power of integrating genetic information with knowledge of the transcriptional regulatory network to identify genes whose biological function is indirectly disrupted by weak genetic associations.

**Figure 3:**
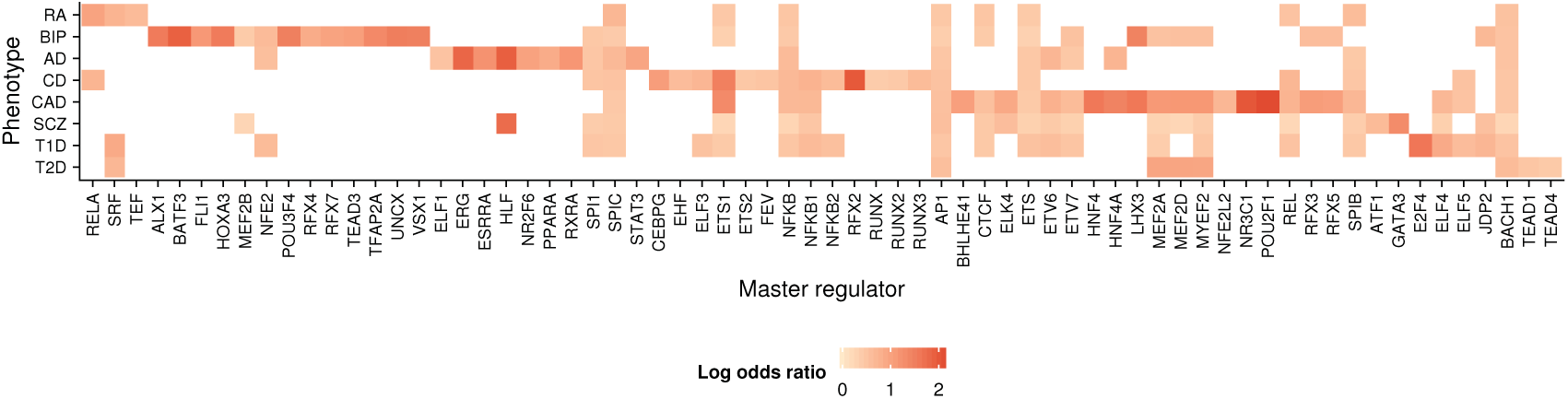
Putative master regulators enriched in enhancer regions harboring weak associations. The maximum enrichment (log odds ratio) is taken for each master regulator over 226 enhancer modules comprising patterns of observed histone modification across 111 reference epigenomes. Only log odds ratios for master regulators with significant enrichment (Fisher’s exact test, BH *q* < 0.05) are shown. Phenotypes are represented by a vector of log odds ratios over each of the master regulators and ordered by hierarchical clustering.

Several of the putative master regulators play known roles in related phenotypes, giving orthogonal evidence for their importance in the eight diseases we studied. We identified *RFX4* as a master regulator in BIP; *RFX4* regulates circadian rhythm, which is disrupted in BIP^30^. We identified *ERG, RXRA*, and *STAT3* in AD. *ERG* mediates AD-like neurodegeneration in Down’s syndrome^31^; *RXRA* alters brain cholesterol metabolism in AD^32^; and *STAT3* mediates amyloid-*β*-induced apoptosis, the classical hallmark of AD^33^. We identified *ELF3* in CD, which is over-expressed in ulcerative colitis (UC) cases^34^, supporting prior work suggesting CD and UC share common genetic factors^35^. We identified *MEF2A* in CAD, which has been previously identified in linkage studies of autosomal dominant CAD^36^.

Additionally, several of the remaining putative master regulators have known biological functions which are *a priori* relevant to the disease they were identified in. We identified *REL* and *ETS1* in multiple diseases, which are known to play a role in immune response^37,38^. We identified *SPI1* in AD, consistent with prior work showing an immune basis for AD^39^. We identified *GATA3* in SCZ and *UNCX* and *TFAP2A* in BIP, which are known to play roles in brain development^40–42^.

We then examined the enhancer regions bound by these master regulators and identified a large number of putative co-factors whose binding sites are directly disrupted by weak associations (**Fig.4**). Moreover, we found that the identified co-factors are specific to both the master regulator and the disease, offering an explanation for how master regulators can be shared between very different diseases. For example, although *NFKB* is enriched in enhancers associated with AD, BIP, CAD, CD, and SCZ, we found that its motif co-occurs with motifs for e.g. *AP1* in AD, *HOX* genes in BIP, *HIC1* in CAD, *IRF3* in CD, and *SP1* in SCZ.

**Figure 4:**
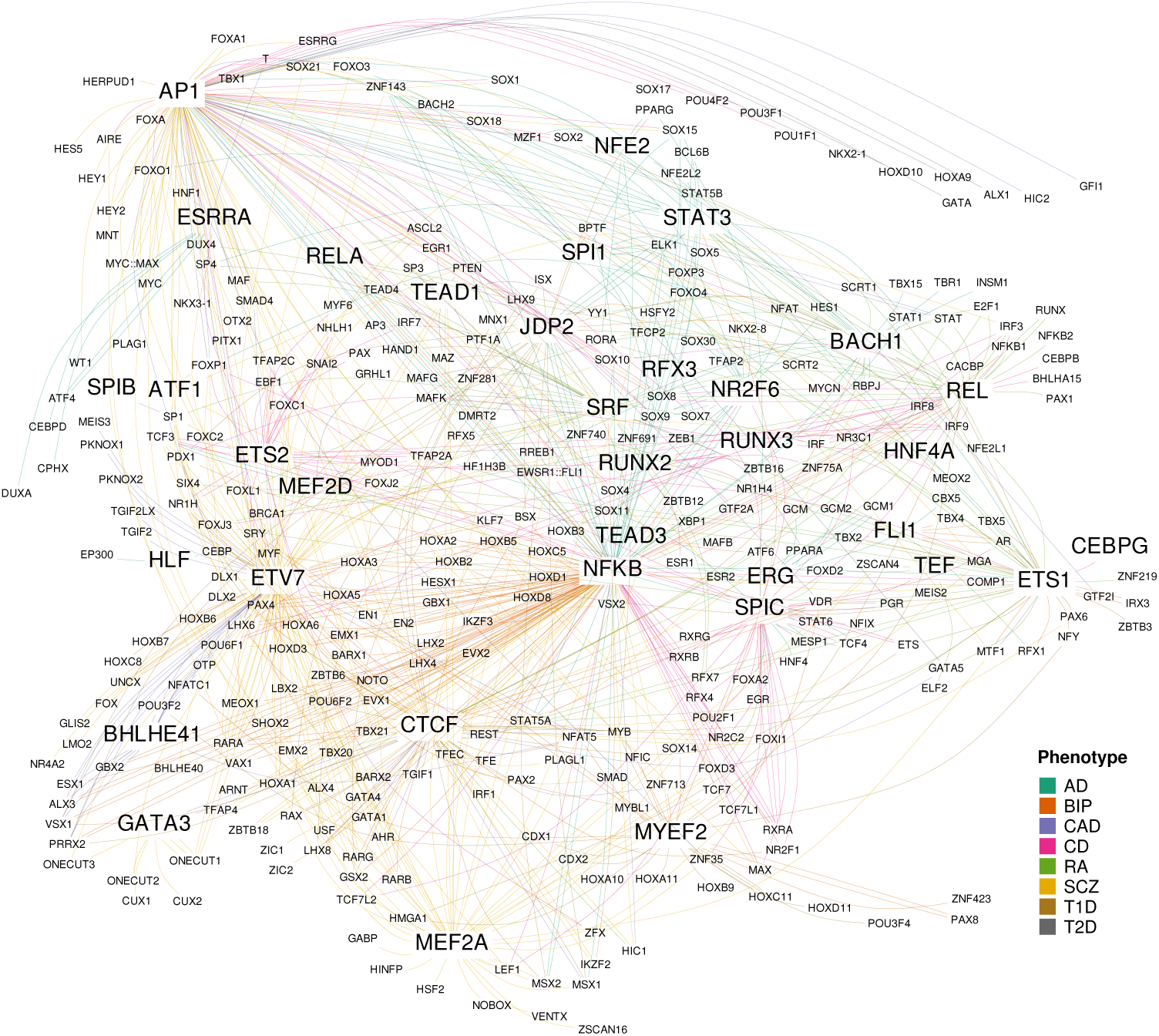
Indirect disruptions of master regulators enriched in enhancer regions by weak associations across eight diseases. Master regulator gene names are given in larger text compared to co-factor gene names. Edges connect master regulators to co-factors for which a motif instance overlaps a weakly associated SNP in an enriched enhancer region and are colored by the associated phenotype. Edges are collapsed such that each interaction appears at most once.

We note that we identified many master regulators in constitutive enhancers (**Supplementary Fig. 5**). One explanation for this result is that we are under-powered to find master regulators in other enhancer modules which cover less of the genome and overlap fewer well-imputed variants. However, even allowing for lack of power in other tissue-specific modules, enrichment in constitutive enhancers runs counter to the hypothesis that different cell type–specific regulators are disrupted in different complex diseases. We hypothesized that although the enhancers might be constitutively marked, the transcription factors which bind to those enhancers would show cell type–specific patterns of expression, explaining their disease specificity. We used RNA-Seq data across 57 reference epigenomes to study the expression of putative master regulators discovered in constitutively marked enhancers and found that indeed they showed diverse patterns of expression (**Supplementary Fig. 6**). For example, *REL, SPI1*, and *ETS1* are predominantly expressed in T cells, consistent with their known tissue-specific functions.

Our results highlight a key distinction between constitutive marking of enhancer-like regions and constitutive activity of distal regulators. However, we found only few master regulators predicted for any disease are clearly expressed in only relevant cell types, possibly due to incomplete profiling of expression across tissues and developmental time points.

## Discussion

In this study, we developed methods to study the role of weak, non-coding variants in complex traits by computing enrichments of weak associations in functional annotations, identifying and correcting for a number of confounders. Across eight complex diseases, we identified relevant regulatory annotations and showed that putative regulatory regions harboring weak associations target relevant downstream genes and are regulated by relevant upstream master regulators. We found enrichments through thousands of independent loci, inviting criticism that in the limit of infinite sample size GWAS will implicate the entire genome^43^. However, we found that in aggregate these thousands of independent loci recurrently disrupt only a small number of pathways, suggesting that improving knowledge of the transcriptional regulatory network offers a way forward in interpreting GWAS.

Our methodology and results highlight two important distinctions in the use of regulatory annotations to identify and re-prioritize weak associations. First, regions marked by enhancer-associated histone modifications are not necessarily active distal regulators. Here, we attempted to characterize putative enhancers by linking them to downstream genes and upstream transcription factors. Second, regulatory annotations predicted on individual reference epigenomes confound constitutive and tissue-specific marking (and activity) of regulatory regions. We showed that A:-means clustering of regulatory regions could deconvolve patterns of histone modification across 111 cell types and tissues, and that measured expression of predicted upstream regulators could deconvolve enhancer activity across 57 cell types.

Our study has several limitations which should be addressed in future work. Most importantly, our methodology finds excesses of associations and motifs in specific annotations and pathways but does not naturally provide measures of confidence for particular loci, genes, or master regulators. We used the BH procedure with *q* = 0.05 throughout to control the false discovery rate (FDR) of rejected hypotheses by setting a new *p*-value threshold; however, this procedure does not estimate an FDR for each hypothesis. In theory, we could use Empirical Bayes to estimate the FDR of each hypothesis^44^. However, our study is arranged as a hierarchy of hypotheses, where the BH procedure is used to screen first-level hypotheses (enhancer modules), and only those rejected are taken forward to test second-level hypotheses (motifs, pathways). Therefore, a novel model would be required to estimate local FDR in our setting, which is beyond the scope of this study. Recent theory proves that applying the BH procedure in this setting does control the FDR over the entire tree of hypotheses^45^; however, the actual FDR over all rejected hypotheses (at any level of the tree) is bounded above by 0.144. Thus, our results should be interpreted as identifying putative enhancer regions, genes, and transcription factors whose role in disease mechanism needs to be confirmed by experimental followup.

We used a heuristic to find a number of of relevant loci and attempted to identify enriched annotations, genes, and regulators without explicitly imposing parametric assumptions about the disease model or causal cell types. However, a number of Bayesian parametric approaches have successfully performed several of these inference tasks^13,46,47^. In particular, the use of spike-and-slab priors allows posterior inferences about the number of causal loci, and the use of regulatory annotations as priors on hyperparameters allow posterior inferences about the importance of different annotations. Importantly, these approaches are either limited to inference on one annotation at a time or do not account for correlation or overlap between related annotations. Alternatively, the structure of the problem naturally suggests a Bayesian network connecting SNPs, enhancers, target genes, and transcription factors; however, such a network directly encodes the transcriptional regulatory network and must somehow account for tissue-specific differences in the network. Further work is needed to combine these ideas and perform more rigorous statistical inference on larger scale data.

Our method is unbiased in the sense that we consider all annotations without imposing any prior information; however, the panel of 127 reference epigenomes we used is itself biased in representation of tissues, leading to several issues. First, we found unexpected enrichments for mucosa cell types across a number of the diseases studied which could be explained by epigenomic similarity to relevant endothelial cell types which were not directly profiled. However, testing this hypothesis will require epigenomic profiling of additional cell types. Second, our definition of a constitutively marked enhancer depends on the proportion of reference epigenomes which the enhancer is annotated by an associated chromatin state. Blood cell types make up a large proportion of reference epigenomes considered here, and therefore putative constitutive regions might not actually be constitutive (leaving aside the distinction between enhancer marks and enhancer activity). Third, enhancer modules in lineages other than blood are smaller than either constitutive or blood-specific modules, making it more difficult to find significant enrichments for these annotations.

More broadly, our methods use annotations of regulatory regions, genes, pathways, and transcription factor binding sites produced by a number of published computational pipelines. These annotations could be sensitive to choices of thresholds and filtering used in each of the pipelines, and therefore our results could also be sensitive to such choices. We took conservative choices in the design of our computational pipeline with regards to correcting for LD and other confounders. However, further work will be needed to characterize the error rates in regulatory annotations and the impact of errors on downstream analyses.

Although we analyzed several million well-imputed variants in each of the eight diseases, we also used finer resolution, higher confidence predictions of regulatory regions, making it more difficult to find significant enrichments. Moreover, although we initially found thousands of loci, they implicate only hundreds of putative enhancer regions of which only a fraction either harbor an enriched motif and or target a gene in an enriched pathway. Future work will need to use more comprehensive panels of variants, better predictions of transcription factor binding sites, and better predictions of distal targets to increase the number of high-confidence testable hypotheses to carry forward to experimental followup.

**Supplementary Information** is linked to the online version of the paper at www.nature.com/nature.

## Acknowledgments

We thank Wouter Meuleman for clustering regulatory regions. We thank Pouya Kheradpour for assistance with the motif analysis pipeline. We thank David Golan, Alexander Gusev, Eric Lander, Alkes Price, Gerald Quon and Zhizhuo Zhang for helpful discussions. A.K.S is supported by an NSF Graduate Research Fellowship (grant #1122374). L.D.W and M.K. are supported by NIH R01HG004037 and R01HG004037-S1.

This study makes use of data generated by the Wellcome Trust Case-Control Consortium. A full list of the investigators who contributed to the generation of the data is available from www.wtccc.orq.uk. Funding for the project was provided by the Wellcome Trust under award 076113 and 085475.

We thank the International Genomics of Alzheimer’s Project (IGAP) for providing summary results data for these analyses. The investigators within IGAP contributed to the design and implementation of IGAP and/or provided data but did not participate in analysis or writing of this report. IGAP was made possible by the generous participation of the control subjects, the patients, and their families. The i–Select chips was funded by the French National Foundation on Alzheimer’s disease and related disorders. EADI was supported by the LABEX (laboratory of excellence program investment for the future) DISTALZ grant, Inserm, Institut Pasteur de Lille, Université de Lille 2 and the Lille University Hospital. GERAD was supported by the Medical Research Council (Grant n° 503480), Alzheimer’s Research UK (Grant n° 503176), the Wellcome Trust (Grant n° 082604/2/07/Z) and German Federal Ministry of Education and Research (BMBF): Competence Network Dementia (CND) grant n° 01GI0102, 01GI0711, 01GI0420. CHARGE was partly supported by the NIH/NIA grant R01 AG033193 and the NIA AG081220 and AGES contract N01–AG–12100, the NHLBI grant R01 HL105756, the Icelandic Heart Association, and the Erasmus Medical Center and Erasmus University. ADGC was supported by the NIH/NIA grants: U01 AG032984, U24 AG021886, U01 AG016976, and the Alzheimer’s Association grant ADGC–10–196728.

## Author Contributions

A.K.S. and L.D.W. developed the methods. A.K.S performed the analysis. A.K.S. and M.K. prepared the manuscript.

## Author information

Reprints and permissions information is available at www.nature.com/reprints The authors declare no competing financial interests.

Correspondence and requests for materials should be addressed to

## Online Methods

### Genome-wide association summary statistics and regulatory annotations

We downloaded summary statistics for AD from the International Genomics of Alzheimer’s Project (see URLs); BIP and SCZ from the Psychiatric Genetics Consortium; CAD from the CARDIOGRAM consortium; CD from the International Inflammatory Bowel Disease Genetics Consortium; RA (https://www.broadinstitute.org/ftp/pub/rheumatoid_arthritis/Stahl_etal_2010NG/RA_GWASmeta2_20090505-results.txt); T1D from the Type 1 Diabetes Genetics Consortium through T1DBase^48^; and T2D from the DIAGRAM Consortium.

International Genomics of Alzheimer’s Project (IGAP) is a large two-stage study based upon genome-wide association studies (GWAS) on individuals of European ancestry. In stage 1, IGAP used genotyped and imputed data on 7,055,881 single nucleotide polymorphisms (SNPs) to meta-analyze four previously-published GWAS datasets consisting of 17,008 Alzheimer’s disease cases and 37,154 controls (The European Alzheimer’s disease Initiative – EADI the Alzheimer Disease Genetics Consortium – ADGC The Cohorts for Heart and Aging Research in Genomic Epidemiology consortium – CHARGE The Genetic and Environmental Risk in AD consortium – GERAD). In stage 2, 11,632 SNPs were genotyped and tested for association in an independent set of 8,572 Alzheimer’s disease cases and 11,312 controls. Finally, a meta-analysis was performed combining results from stages 1 & 2.

We downloaded ChromHMM segmentations from the Roadmap Epigenomics project; clustered regulatory regions from the Regulatory Regions Map; genic annotations from the GENCODE project; CAGE–predicted transcription start sites (ftp://genome.crg.es/pub/Encode/data_analysis/TSS/Gencodev10_CAGE_TSS_clusters_May2012.gff.gz); predicted motif instances from the ENCODE project; and motif enrichments (predicted regulators) from the Roadmap Epigenomics project.

### Imputation of summary statistics

We downloaded Thousand Genomes (1KG) reference haplotypes in OXSTATS format (September 2013 version, no singletons). We used ImpG-Summary with default parameters and all 1KG samples to impute summary statistics for BIP, CAD, RA, T1D, and T2D into all SNPs with MAF > 0.01 in 1KG European samples.

In order to assign signs of effects for T1D (for which odds ratios were not published), we imputed genotypes for the Wellcome Trust Case Control Consortium study of T1D and took the sign from the single-SNP association test.

We downloaded probe identifiers, hg19 positions, and strand information (http://www.well.ox.ac.uk/~wrayner/strand/) to convert positions to hg19 and used GTOOL version 0.7.5 to align all genotypes. We used PLINK version 1.09b to produce hard genotype calls with genotype probability threshold 0.99 and remove all SNPs and samples excluded from the original study. We used SHAPEIT2 v2.r644 (ref.^49^) to exclude unalignable SNPs and phase the case and control cohorts independently for each autosome. We used default values for all model parameters.

We used IMPUTE2 version 2.3.0 (ref.^50^) to impute into all SNPs and indels with MAF in European samples > 0.01. We divided the autosomes into 5 MB windows and threw out windows with fewer than 100 array probes. We used SNPTEST version 2.5.1 (ref.^51^) to compute association *β*-values using maximum likelihood estimates of an additive model. We included 10 principal components computed using GCTA 1.24 (ref.^52^) on the hard-called array genotypes. We made extensive use of GNU parallel^53^ to facilitate the analysis.

### Visualization of functional enrichment

For each disease and annotation, we compared the observed number of overlaps with the annotation against the expected number of overlaps at each rank threshold (every 1,000 SNPs). Given *K* of *N* total variants overlap a functional region, the expected number of overlaps in the top *n* variants is *K* × *n*/*N*. We plotted the difference normalized by the total number of overlaps genome-wide. We used BEDTools version 2.24 (ref.^54^) to compute overlaps.

To pick an empirical *p*-value cutoff, we first computed the convex hull of each curve, then computed the elbow point as the first inflection point in the convex hull. To compute inflection points, we approximated the second derivative of the curves by twice taking the difference of adjacent points normalized by the interval size and took the first point where the second derivative changed sign. We took the least stringent *p*-value cutoff (maximum elbow point) to be the empirical cutoff to carry forward in the analysis.

### Statistical test for functional enrichment

For each disease and annotation, we applied a one-sided permutation test comparing the observed count of variants in the annotation meeting the new *p*-value cutoff against the null distribution of the analogous counts over 10,000 resampled sets. We resampled variants with replacement (to reduce memory usage) from outside the regions of interest and matched on number of LD partners (*r*^2^ > 0.1), minor allele frequency (in bins of width 0.05), and distance to nearest transcription start site (rounded to the nearest kilobase).

We computed *p*-values by counting the number of resampled sets with at least as many overlaps as the original data. We used the Anderson-Darling test to test whether the null distribution was approximately Gaussian. We reported *z*-scores based on the mean and variance of count of overlaps over the resampled sets. We applied the Benjamini–Hochberg procedure with *q* = 0.05 to control the false discovery rate.

### Controlling for LD

We computed pairwise correlations between pairs of variants in the Thousand Genomes European samples within 1 megabase and with *r*^2^ > 0.1. We pruned to a desired threshold by iteratively picking the top-scoring variant (breaking ties arbitrarily) and removing the tagged variants until no variants remained.

To adapt our visualization to account for LD between weak associations, we first pruned the list of imputed variants according to reference LD information to obtain a list of pairwise independent tag variants with *r*^2^ < 0.1. We then modified the above formulas by summing a fractional haplotype score over loci instead of counting variants, defined as the proportion of variants in the locus falling in a functional region. Then, at each rank threshold we compared the observed total score against the expected score. The expected score was computed as the total genome-wide score multiplied by the proportion of loci meeting the rank threshold, and the difference was normalized by the total score genome-wide.

### Pathway enrichment of enhancer targets

We used GREAT to test for enrichment of enhancer regions in gene pathways. For each enhancer module, we defined the foreground as the set of regions containing associated SNPs meeting the empirical *p*-value cutoff and the background as all regions in the module.

We used Phenotype-Genotype Integrator to retrieve a list of known genes for each disease and matched linked genes in each enriched pathway to known genes based on gene names.

To prune enriched pathways, we downloaded the basic version of Gene Ontology in Open Biomedical Ontologies format and built the specified directed acyclic graph connecting terms to their parents. We performed depth-first traversal of the graph starting from enriched terms and took nodes which were never reached from a child node as the most specific enriched terms.

### Motif enrichment of upstream regulators

For each enhancer module, we first filtered motifs based on sequence enrichment as previously described^55^.

For each combination of disease, module, and sequence-enriched motif, we constructed a 2 × 2 contingency table counting enhancer regions partitioned by presence of the motif and orthogonally by presence of a weak association (based on our empirical *p*-value cutoff). We restricted the set of regions to the domain on which motifs were discovered (excluding coding regions, 3’ UTRs, transposons, and repetitive regions) and additionally to the subset of regions which harbor an imputed SNP for the disease. We computed one-sided *p*-values using Fisher’s exact test.

For each putative master regulator, we re-scanned regions containing both a motif instance and a weak association for any motif instances overlapping the associated SNP. We used manual annotation of the motifs to collapse motifs by transcription factor.

We used the transcription factor gene names to visualize expression of the upstream regulators across 57 reference epigenomes. We normalized the expression RPKM by scaling the maximum value to 1 in order to put expression of each TF on the same scale.

### Code availability

Code used to perform the analysis is available from https://www.github.com/aksarkar/frea and https://www.github.com/aksarkar/frea-pipeline

### URLs

- International Genetics of Alzheimer’s Project http://web.pasteur-lille.fr/en/recherche/u744/igap/igap_download.php
- Psychiatric Genetics Consortium http://www.med.unc.edu/pgc
- Coronary Artery Disease Genome-wide Replication and Meta-analysis Consortium http://www.cardiogramplusc4d.org/
- International Inflammatory Bowel Disease Genetics Consortium http://www.ibdgenetics.org/
- Diabetes Genetics Replication and Meta-analysis Consortium http://diagram-consortium.org/
- T1DBase http://www.t1dbase.org/
- Rheumatoid arthritis summary statistics https://www.broadinstitute.org/ftp/pub/rheumatoid_arthritis/Stahl_etal_2010NG/
- Thousand Genomes reference data (http://mathgen.stats.ox.ac.uk/impute/)
- Roadmap Epigenomics http://egg2.wustl.edu/roadmap/web_portal/
- Regulatory Regions Map https://www.broadinstitute.org/~meuleman/reg2map/
- GENCODE version 10 ftp://ftp.sanger.ac.uk/pub/gencode/Gencode_human/release_10/
- GREAT: Genomic Regions Enrichment of Annotations Tool http://bejerano.stanford.edu/great/public/html/
- Gene Ontology http://geneontology.org/
- Phenotype-Genotype Integrator https://www.ncbi.nlm.nih.gov/gap/phegeni
- ENCODE motifs http://compbio.mit.edu/encode-motifs/

